# Bacteria-mimetic bioadhesives with multivalent mucoadhesion and drug-compatible delivery

**DOI:** 10.64898/2026.04.15.718429

**Authors:** Evan Johnston, Megan Farrow, Zhen Yang, Aram Bahmani, Yin Liu, Xin Huang, Jing Yan, Jianyu Li

**Affiliations:** Department of Mechanical Engineering, McGill University, Montreal, Quebec, H3A 0C3, Canada; Department of Biomedical Engineering, McGill University, Montreal, Quebec, H3A 0C3, Canada; Department of Molecular, Cellular and Developmental Biology, Quantitative Biology Institute, Yale University, New Haven, Connecticut 06520, United States

## Abstract

The ability to adhere to mucus-lined tissues underpins a range of biomedical devices and therapies. However, many existing strategies rely on covalent bonding chemistries and can be unstable, cytotoxic, or incompatible with therapeutics. Here, we present a bacteria-mimetic bioadhesion strategy inspired by *Vibrio cholerae*. A short Bap1-derived adhesion peptide is grafted onto chitosan to strengthen mucus interactions through multivalent, cooperative secondary bonding, while preserving pH-triggered interfacial bridging behavior. Bacterial peptide grafting significantly increases adhesion energy on porcine intestine, and when paired with a tough hydrogel matrix achieves adhesion energies >400 J/m^2^ without forming covalent bonds to tissue. Confocal imaging reveals deep tissue penetration (∼80 μm) with markedly enhanced mucin binding and no loss of cytocompatibility. Ex vivo intestinal delivery and in vitro drug release tests demonstrate improved drug transport and tissue exposure compared to carbodiimide-mediated covalent bonding strategy. These findings establish a bacteria-mimetic bioadhesion strategy for tissue repair and drug delivery.

**Novelty Statement:** Bioadhesive designs have drawn inspiration from nature such as mussel-inspired catechol chemistry and gecko-inspired dry adhesion. In contrast, bacterial adhesion mechanisms, despite enabling robust colonization of mucus-lined tissues under demanding conditions, remain largely overlooked. This work introduces a bacteria-mimetic bioadhesion strategy that selectively repurposes a short, non-pathogenic peptide derived from a *Vibrio cholerae* adhesin to enhance bioadhesion through multivalent, cooperative physical interactions rather than covalent bonding. This strategy significantly toughens adhesion even on chitosan, a polymer already rich in adhesive functional groups. By decoupling bacterial adhesion function from pathogenic risk, this study establishes bacterial adhesion peptides as a safe, modular, and mechanistically distinct foundation for next generation bioadhesives with improved drug compatibility.

## Introduction

Bioadhesives play a critical role in enabling medical devices, tissue repair strategies, and drug delivery systems to function effectively in the human body^1, 2^. Among these applications, adhesion to mucus-lined tissues is important, as mucus-covered interfaces are found throughout gastrointestinal, respiratory, ocular, and reproductive systems. Adhesion on these tissues is inherently challenging due to constant hydration, continuous mucus renewal, and dynamic mechanical environment^3, 4^. Although mucoadhesive polymers such as chitosan^5^, carbomers^6^, and cellulose derivatives^7^ have been reported, they still suffer from limited adhesion strength, reduced stability under physiological conditions, and potential cytotoxicity at higher doses or with prolonged exposure^8, 9^. To improve adhesion, covalent bonding strategies have been introduced, enabling permanent attachment to mucosal tissues. However, such approaches sacrifice reversibility and may induce local inflammation or cytotoxicity. Moreover, in drug delivery contexts, reactive coupling chemistries can immobilize therapeutics or interfere with their transport, limiting efficacy in chronic or repeated-use applications.

In contrast to synthetic approaches, many biological systems can adhere strongly to wet, dynamic biological surfaces using non-covalent yet highly effective mechanisms^10, 11^. For instance, bacteria, including pathogenic species like *Vibrio cholerae*, are able to colonize mucosal surfaces despite the dynamic environment of the gastrointestinal tract^12^. This capability is enabled by specialized adhesion proteins that engage mucosal glycans and lipid membranes through multivalent secondary interactions, including hydrogen bonding, electrostatic interactions, and CH-π interactions^13, 14^. Harnessing these bacterial adhesion principles offers an attractive alternative for designing strong yet drug-compatible mucoadhesive materials^15^.

Recent progress has identified these specific protein motifs that underpin bacterial adhesion on intestine tissues. In *Vibrio cholerae*, the biofilm-associated protein 1 (Bap1) plays a central role in surface attachment^16, 17^, with structural studies revealing lectin-like interactions between Bap1 domains and anionic polysaccharides^18^. Notably, a 57-amino-acid adhesive region within Bap1 displays a dual-binding mechanism, in which a central aromatic-rich segment anchors to lipid bilayers, while peripheral repeating motifs, including the WKTKTVPY sequence, enhance bioadhesion through multivalent avidity effects, thereby engaging both lipid membranes and glycan-rich substrates (Figure 1A)^13^. These findings suggest that bacterial adhesion principles may be distilled into short, synthetic peptide motifs capable of enhancing interfacial affinity without introducing pathogenic risk.

**Fig. 1.**
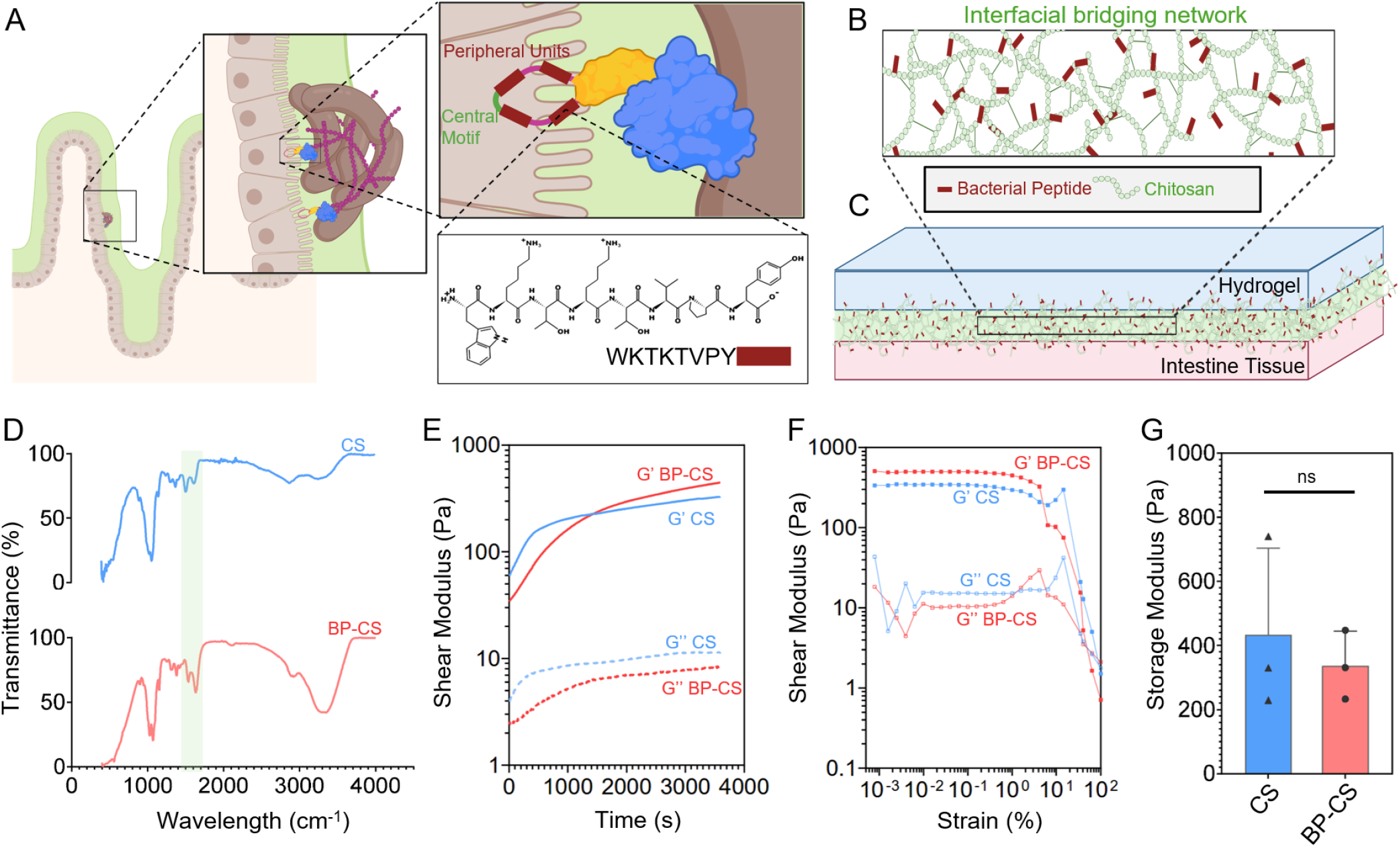
Design and characterization of bacterial peptide conjugated chitosan reliant on bacterial adhesion principles. (A) Schematic representation of repeating peripheral WKTKTVPY sequence in the dBap1 adhesin. Schematic representation of (B) bacterial peptide modified chitosan (BP-CS) and its application as a (C) bridging polymer. (D) FTIR spectra for both CS and BP-CS. Increased absorbance at around 1650cm^−1^ correspond to C=O stretch, indicating successful modification. (E) Time-dependent rheological measurements of storage modulus (G’) and loss modulus (G”) for pristine CS and BP-CS. (F) Strain sweep rheology demonstrates the mechanical stability of both formulations, with failure around 10% strain. (G) Plateau storage modulus for CS and BP-CS. Rheological analysis profiles are comparable between CS and BP-CS in all cases. Data are presented as means ± standard deviation, (ns p>0.05).

Here, we translate this concept into a bacteria-mimetic bioadhesive design by grafting a short Bap1-derived adhesion peptide (WKTKTVPY) onto chitosan, a widely studied mucoadhesive polymer. Under mildly acidic conditions, protonated chitosan electrostatically interacts with negatively charged components of mucus^5, 19^, and can form pH-responsive bridging networks between hydrogels and tissues for bioadhesion^20-22^. By grafting the Bap1-derived bacterial peptide onto chitosan to create bacterial peptide-modified chitosan (BP-CS), we aim to selectively enhance interfacial affinity and bioadhesion through multivalent, non-covalent interactions. We hypothesize that the combination of *V. cholerae*-derived adhesion and pH-triggered gelation behavior of chitosan align well with applications within the mildly acidic environment of large intestine and colon environment (pH 6-7)^23, 24^.

In this study, we test the adhesion performance of BP-CS between porcine intestine with either single-network or double-network hydrogels. Further, we combine rheological tests and confocal imaging to probe the adhesion mechanism of BP-CS on intestinal tissue. Finally, we assess cytocompatibility and intestinal drug delivery to determine whether non-covalent adhesion can maintain macromolecular delivery relative to covalent bonding strategies. Together, this work establishes bacterial adhesion principles as a safe and effective strategy for engineering bioadhesive systems.

## Results and discussion

### Design principle and polymer modification

The adhesion mechanism of BP-CS is designed to couple chitosan bridging network formation with peptide-mediated interfacial interactions (Figure 1B). BP-CS is applied under acidic conditions, where chitosan remains soluble and mobile, allowing the polymer chains to diffuse and penetrate both local intestinal tissue and the hydrogel matrix. Triggered by the neutral pH at the tissue–hydrogel interface, chitosan subsequently undergoes self-association and physical crosslinking between polymer chains, leading to the formation of a chitosan bridging network spanning the interface^**25**^. Concurrently, the grafted Bap1-derived peptide motifs promote multivalent physical interactions with the intestinal mucus layer, enforcing adhesion through non-covalent interactions (Figure 1C).

Successful conjugation of the bacterial peptide to CS was confirmed through Fourier-transform infrared (FTIR) spectroscopy and nuclear magnetic resonance (NMR) spectroscopy. FTIR spectra revealed characteristic peaks corresponding to amide bond formation, indicating successful coupling of the peptide to the CS backbone. Specifically, an increase in absorbance at approximately 1650 cm^−1^, corresponding to C=O stretching in amide bonds, was observed in BP-CS compared to unmodified CS (Figure 1D). NMR analysis further supported these findings, with new signals appearing in the BP-CS spectrum, consistent with the chemical structure of the conjugated peptide (supplemental Figure S1). Notably, chemical signal shifts around 3–4 ppm were observed, likely associated with aliphatic side chains or modifications to the CS backbone caused by peptide attachment.

### Preservation of pH-triggered gelation at interface

As pH-induced chitosan gelation is essential for forming interfacial bridging network, we next examined whether peptide conjugation alters gelation and mechanical behavior. Gelation was induced by sodium bicarbonate in dissolved chitosan solutions^26^, and rheological measurements were conducted to compare pristine CS and BP-CS. Time-sweep tests revealed similar evolution of storage (G′) and loss (G″) moduli for CS and BP-CS following pH trigger by NaHCO_3_ addition (Figure 1E), indicating that peptide conjugation does not alter gelation kinetics. For mechanical responses, strain sweep measurements showed comparable shear moduli in linear viscoelastic regions, as well as similar critical strains for softening (Figure 1F). Although BP-CS exhibited a slightly reduced plateau storage modulus, this difference was not statistically significant (Figure 1G). The comparable mechanical responses were also seen in frequency sweep tests across a wide range of frequency (supplemental Figure S2). Collectively, these results demonstrate that grafting of the Bap1-derived bacterial peptide onto chitosan preserves its pH-triggered gelation behavior and mechanical properties.

This preservation of gelation and mechanical behavior is mechanistically consistent with the chemical composition of the grafted WKTKTVPY peptide sequence^27^. Notably, the sequence contains two lysine residues, but no carboxylic acid–bearing residues. Therefore, peptide addition could reinforce the local positive charge density, while avoiding the formation of unwanted electrostatic complexes that could otherwise disrupt pH sensitivity or induce polyelectrolyte complexation. Rather, the majority of amino acids in this sequence are neutral or weakly positive charged at neutral pH, favoring hydrophobic associations and electrostatic attraction with mucin.

### Enhanced adhesion to intestine tissue

After confirming preserved chitosan gelation and mechanical behaviors, we next tested whether BP-CS, presenting multivalent bacterial adhesion motifs, could further enhance adhesion between hydrogels and intestine tissue. By engaging multiple binding sites simultaneously, addition of bacterial derived peptide is expected to promote interactions with biological membranes and abiotic surfaces, resulting in more robust and durable adhesion under physiological conditions. Adhesion performance was quantified with adhesion energy, measured using T-peel tests on ex vivo porcine lower intestine (pH ∼7), enabling both mucosal interaction and chitosan gelation (Figure 2A). CS or BP-CS was applied as a bridging polymer between intestinal tissue and either single-network PAAm hydrogels or double-network Alg–PAAm hydrogels.

**Fig. 2.**
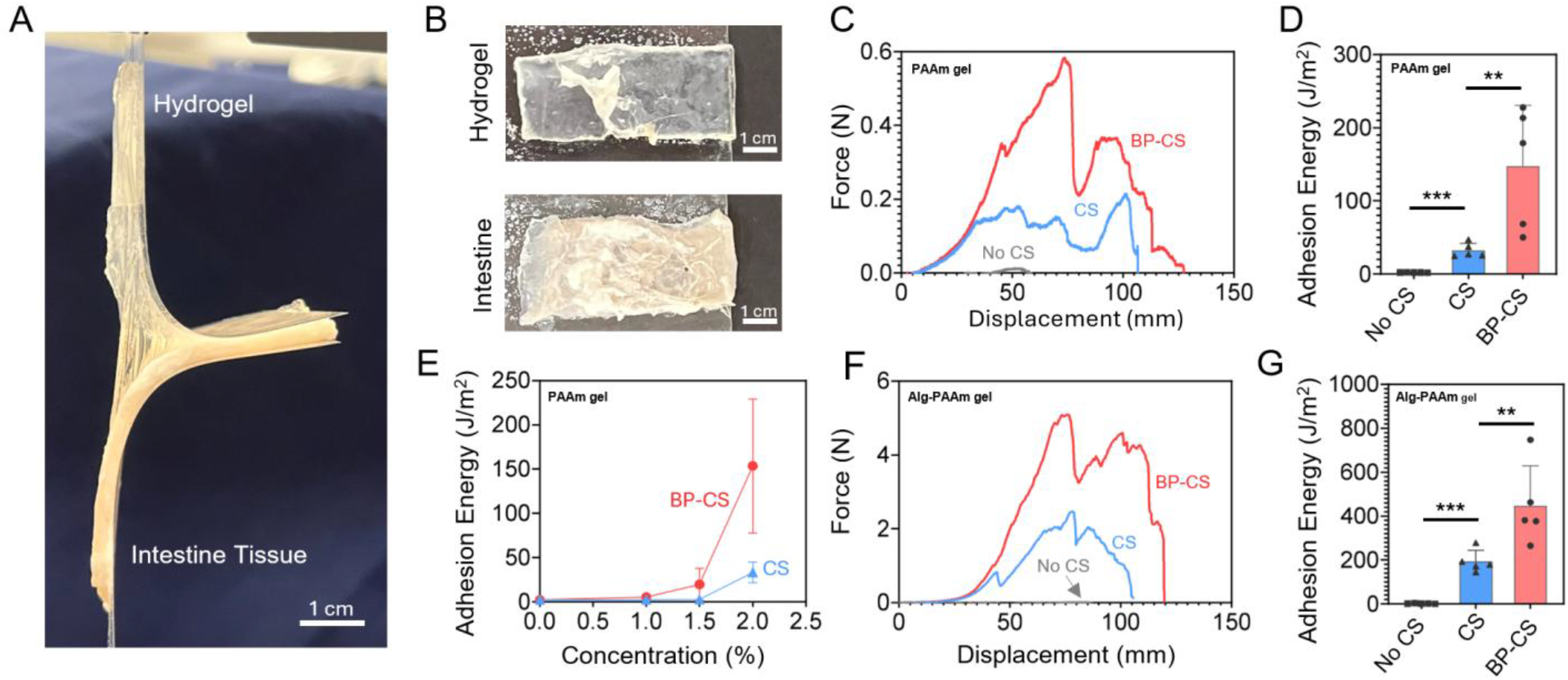
Adhesion testing results reveal enhanced adhesion energy for BP-CS adhesive system. (A) T-peel testing setup for evaluating adhesion energy of material system on ex vivo porcine large intestine. (B) Representative fractured samples indicate some adhesive failure and mixed fracture occurs, where residual tissue may remain attached to hydrogel. (C) Force-displacement curves and (D) adhesive performance of intestine tissue adhered to single-network PAAm gels using 2% pristine CS or BP-CS. (E) Adhesive performance for PAAm hydrogel with various CS or BP-CS concentrations (0, 0.5, 1.0, 2.0%). Adhesion was only established with 2% pristine CS or with at least 1.5% BP-CS. (F) Force-displacement curves and (G) adhesive performance of intestine tissue adhered to tough Alg-PAAm gels using 2% pristine CS or BP-CS. Data are presented as means ± standard deviation (***p < 0.001, **p < 0.01).

PAAm hydrogels exhibit minimal hysteresis and were therefore selected to provide a low-background platform, whereby measured peel energy is dominated by the interfacial bonding of the chitosan bridging layer^28^. Effectively, this design allows us to more cleanly isolate the contribution of the Bap1-derived multivalent motifs to bioadhesion. Further, PAAm gels were selected as an adhesive matrix due to their deformable and highly compliant properties, which allow them to form conformal contact with uneven or delicate surfaces like intestinal tissue. This soft, adaptable nature makes PAAm gels an excellent model for mimicking the mechanical behavior of soft tissues and evaluating the adhesive performance of mucoadhesive systems under physiologically relevant conditions. In contrast, Alg– PAAm double-network gels were used as a tougher, energy-dissipating matrix that adds substantial bulk dissipation and toughening during peeling. These tough gels combine the elasticity of PAAm with the ionic crosslinking of alginate, providing a mechanically robust platform that can dissipate stress during peeling and simulate more challenging adhesive conditions^29, 30^. Use of the double network gel allows us to probe BP-CS performance under higher mechanical loads and to assess the upper bound/limit of adhesion achievable when interfacial binding is paired with a tough hydrogel matrix. This system enables verification of the use of BP-CS for applications requiring stronger adhesion and mechanical stability.

For the PAAm gels, force-displacement curves (Figure 2C) reveal that the inclusion of pristine CS alone as an interfacial bridging polymer significantly improves mucosal tissue – gel-adhesion performance. However, use of modified BP-CS in place of CS resulted in substantial enhancement in peel force, as evidenced by greater force observed during peeling. The calculated adhesion energy (Figure 2D) supports these findings, with BP-CS achieving an average adhesion energy of approximately 148 J/m^2^ compared to the 32 J/m^2^ of unmodified CS and negligible adhesion for the no-CS condition.

The enhanced adhesion performance with BP-CS is also evident at lower concentrations (Figure 2E). While pristine CS only established measurable adhesion to intestinal tissue at 2 wt%, BP-CS was able to form detectable adhesion even at a reduced concentration of 1.5 wt%. Although the adhesion energy at this concentration was modest (∼20 J/m^2^), this result underscores that peptide grafting enables adhesion under conditions where unmodified CS is ineffective. This behavior likely reflects enhanced peptide–tissue interactions, potentially arising from increased local cationic charge density and multivalent secondary bonding introduced by the grafted peptide motifs.

To assess whether this adhesion enhancement could be further amplified with a tough hydrogel matrix, we also evaluated BP-CS in combination with double-network Alg–PAAm hydrogels, which have previously demonstrated to provide bulk dissipation to toughen bioadhesion. Similar to the trends observed with single-network PAAm gels, BP-CS markedly outperformed pristine CS and the no-CS control when paired with Alg–PAAm gels. Force–displacement curves (Figure 2F) revealed substantially higher peeling forces for BP-CS, and the calculated adhesion energy reached an average of ∼446 J/m^2^, more than twice that achieved with pristine CS (∼193 J/m^2^) (Figure 2G).

These results highlight the effectiveness of peptide modification in enhancing bioadhesion across different gel systems. BP-CS increased adhesion energy by factors of approximately 4.6 and 2.3 for single network PAAm and double network Alg–PAAm hydrogel systems, respectively. We attribute these improvements to peptide-mediated interactions with the mucosal interface, including enhanced electrostatic interactions, and hydrophobic associations with mucins and epithelial surfaces.

### Diffusion and mucin binding

To investigate how BP-CS enables adhesion to intestine tissue, we examined both diffusion of chitosan in ex vivo porcine intestine and in vitro mucin binding. For the diffusion study, CS and BP-CS were labeled with FITC and imaged by confocal microscopy to quantify penetration into the intestinal mucosa (Figure 3A). Both CS and BP-CS penetrated the mucosal layer effectively, reaching comparable diffusion depths of ∼80 μm (Figure 3B). This phenomenon can be attributed to the fact that chitosan penetration is mainly governed by its diffusivity and the time scale of pH-driven neutralization/deprotonation that induces formation of a bridging network. In line with our rheological results, BP grafting minimally affected pH-triggered gelation and is small relative to the chitosan backbone, supporting the similar diffusion profiles. Notably, this diffusion depth exceeds the typical thickness of the surface mucin layer, suggesting that the chitosan network can extend into the mucosa to resist rapid mucus turnover and potentially contact epithelial cells and their lipid membranes. Given that the bacterial peptide exhibits lipid-binding activity, such interactions may contribute to the improved adhesion observed for BP-CS.

**Fig. 3.**
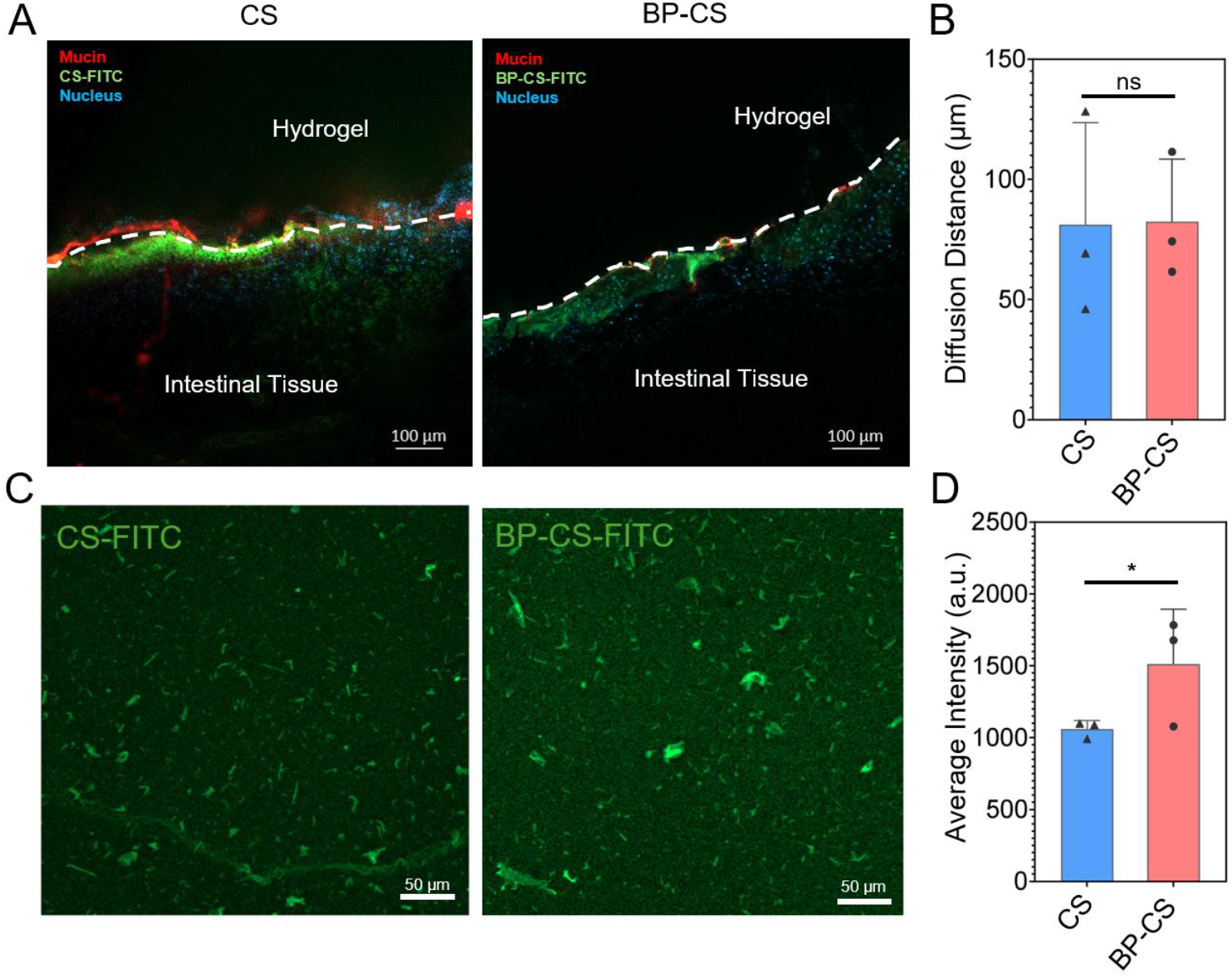
Diffusion into ex vivo tissue and mucin binding comparison between pristine chitosan (CS) and bacteria peptide-modified chitosan (BP-CS). (A) Confocal microscopy images showing the diffusion of FITC-labeled pristine CS and BP-CS into ex vivo porcine intestine tissue, and (B) quantification of diffusion distances confirms that both formulations penetrate similarly into the tissue (approximately 80 μm). (C) Confocal imaging of mucin binding for both CS and BP-CS, quantified by average fluorescence intensity as seen in (D). Data are presented as means ± standard deviation (*p < 0.05, ns p>0.05).

While diffusion and network formation were preserved, we hypothesized that the enhancement in adhesion arises from increased affinity between chitosan and mucin. Mucin is a major component of mucus and mucin binding is often considered a key factor in mucoadhesion^31^. To evaluate this hypothesis, we performed a mucin binding assay adapted from fluorescence-based methods used to quantify bacterial and protein binding to mucin glycoproteins^31^. Mucin-coated wells were treated with FITC-labeled CS or BP-CS, rinsed, and imaged; fluorescence intensity was used as a proxy for chitosan retention (Figure 3C). Our quantification showed a significantly higher fluorescence intensity for BP-CS compared to the pristine counterpart, indicating stronger mucin binding and improved retention. Together with the diffusion result, these findings suggest that adhesin-derived peptide grafting enhances bioadhesion primarily by increasing affinity for mucin glycoproteins.

### Biocompatibility

With the adhesion enhancement by peptide grafting established, we next assessed the cytocompatibility of BP-CS with in vitro cell culture. EDC/NHS-mediated crosslinking has been reported to induce cytotoxicity, particularly at elevated concentrations, due to its non-specific reactivity with biomolecules^32, 33^. Because BP–CS adhesion relies on reversible secondary interactions such as hydrogen bonding and electrostatic interactions, we hypothesized that BP grafting would not compromise cell viability. Affirming this, live/dead fluorescence imaging revealed predominantly viable (green) cells across all four tested conditions, including PAAm alone, BP-CS, PAAm + BP-CS, and control medium (Figure 4A). Cells also maintained normal morphology and uniform distribution. Quantitative viability analysis further confirmed these observations as cell viability exceeded 85% in all groups, with no statistically significant differences among BP-CS, PAAm, or their combined system (Figure 4B). Importantly, incorporation of BP-CS into the interface of PAAm hydrogels did not reduce viability relative to PAAm alone.

**Fig. 4.**
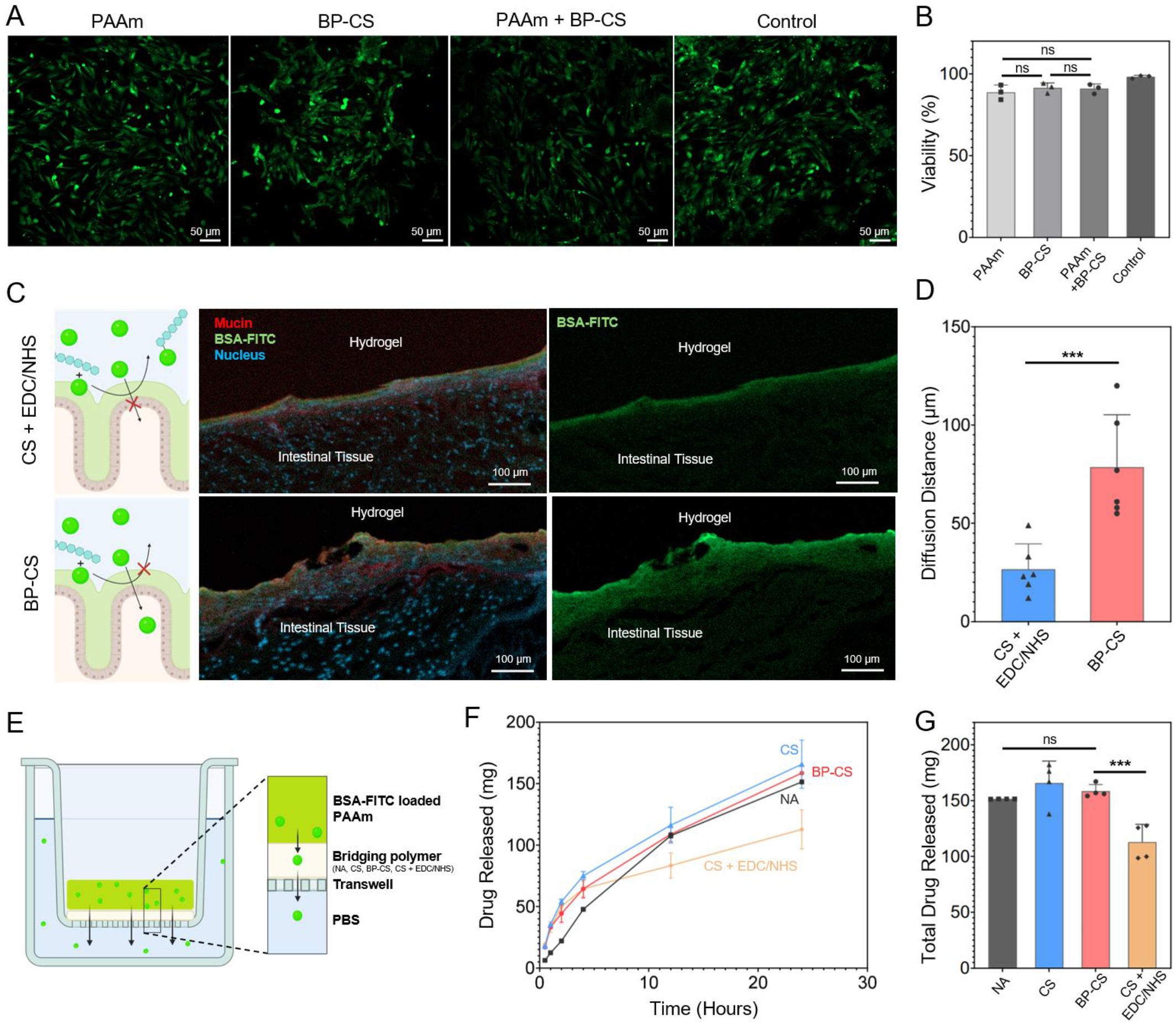
Biocompatibility and diffusion of model drug. (A) Fluorescence microscopy images of HVFF cells exposed to extraction medium prepared with PAAm gel alone, BP-CS, PAAm + BP-CS, or control medium show no significant cytotoxicity. (B) Cell viability assay results confirm high biocompatibilities for all conditions, with viability exceeding 85% across treatments. (C) Confocal microscopy images show the penetration of BSA-FITC into porcine intestinal tissue with BP-CS and CS + EDC/NHS, with a limited diffusion of drug observed for covalent adhesion. The adhesive system was applied to ex vivo porcine intestine tissue (cells stained blue, mucin stained red), and diffusion of model drug, BSA-FITC (green) was evaluated after 6 hours. (D) Quantification of diffusion distances reveals significantly higher penetration for BP-CS compared to CS + EDC/NHS. (E) Schematic of drug diffusion kinetics study demonstrating the use of trans wells, allowing for the diffusion of BSA-FITC loaded in a PAAm gel through the test bridging polymers. Drug release kinetics (F) and total drug release (G) for this study are shown with the use of no bridging polymer (NA), pristine chitosan (CS), bacterial peptide modified chitosan (BP-CS) and chitosan relying on EDC/NHS chemistry (CS + EDC/NHS), Data are presented as means ± standard deviation, (***p < 0.001, **p < 0.01*p < 0.05, ns p > 0.05).

### Drug delivery

As BP-CS strengthens adhesion without covalent bonding, we hypothesized that it could better preserve drug mobility at the adhesive interface. This property is particularly relevant for chronic or repeated-use applications (e.g., inflammatory bowel disease), where reactive crosslinking chemistries may immobilize therapeutics or induce local tissue damage. To test this hypothesis, we loaded the PAAm hydrogel with FITC–BSA as a model macromolecular therapeutic (Figure 4C) and compared intestinal drug delivery using BP-CS versus a covalent-adhesion control (CS + EDC/NHS). FITC-BSA has been widely used to directly visualize and quantify macromolecular diffusion in tissue and also serves as a representative protein therapeutic containing carboxyl and amine groups^34^. The CS + EDC/NHS control was selected due to the frequent use of carbodiimide coupling as a standard covalent bonding strategy used in various tough adhesive systems. This methodology operates by activating carboxyl groups to form amide bonds with amines at the tissue and/or hydrogel interface^26, 30^.

Confocal imaging showed that both conditions (CS + EDC/NHS, and BP-CS) permitted BSA-FITC diffusion into the mucosal layer; however, BP-CS achieved significantly deeper penetration and a more uniform intratissue distribution, with higher fluorescence intensity than the control (Figure 4C). Consistently, quantitative analysis revealed an approximately threefold increase in diffusion depth for BP-CS relative to CS + EDC/NHS (Figure 4D).

To further quantify drug release behavior, we measured drug diffusion kinetics using a Transwell setup (Figure 4E). Drug transport across interfaces containing BP-CS, CS + EDC/NHS, or pristine CS was monitored over time. The release profile of BP-CS closely resembled those of the non-adhesive (NA) and pristine chitosan (CS) conditions (Figure 4F), indicating minimal interaction between BP-CS and the model drug. In contrast, the CS + EDC/NHS condition exhibited a markedly slower release rate and a significantly reduced cumulative drug release over 24 hours (Figure 4G), consistent with drug immobilization at the covalently crosslinked adhesive interface seen in the intestinal drug delivery experiment.

Together, these results demonstrate that BP-CS supports efficient drug diffusion both in terms of release kinetics and tissue exposure. The non-covalent adhesion mechanism of BP-CS, in addition to providing strong adhesion and high cytocompatibility, minimizes off-target crosslinking with protein therapeutics and enables faster, more complete drug release compared to EDC/NHS-mediated covalent strategies. This feature is particularly important for delivering macromolecular therapeutics, such as proteins and nucleic acids, to mucus-lined tissues like the gastrointestinal tract, where rapid mucus turnover and enzymatic degradation limit therapeutic residence time. By combining robust bioadhesion with drug compatibility, peptide-functionalized chitosan represents a promising platform for future translational applications in gastrointestinal repair and drug delivery.

## Conclusions

To conclude, this work presented a bacteria-mimetic bioadhesion strategy inspired by the *Vibrio cholerae* adhesin Bap1. By selectively upgrading interfacial affinity while preserving the pH-triggered gelation behavior of chitosan, this approach enabled strong, non-covalent adhesion on mucus-lined tissues. Compared to covalent bonding strategies, bacterial peptide-grafted chitosan achieved strong tissue adhesion, while avoiding irreversible crosslinking, cytotoxicity, and drug immobilization.

This study primarily evaluates ex vivo adhesion and drug delivery performance. In vivo validation with animal experiments is needed to assess efficacy in physiological conditions, long-term biocompatibility and potential immunogenicity. Future work could also optimize the peptide sequence and density to further enhance adhesion performance. Beyond intestinal tissue, this strategy may be extended to other mucus-lined organs, including oral cavity, respiratory tract, ocular surface, and urogenital tissues. In addition, the bacterial adhesion peptides could be integrated into other polymeric adhesive systems, analogous to the widespread adoption of mussel-inspired catechol chemistry, enabling modular and tissue-specific bioadhesive design.

More broadly, this work highlights the promise of converging bacterial adhesion biology with biomaterials engineering. By learning from how microbes robustly colonize soft, hydrated, and dynamically renewing interfaces, new biomimetic strategies can emerge for designing next generation bioadhesives, opening opportunities across drug delivery, tissue repair, and biomedical device development.

## Experimental section

### Synthesis of peptide-modified chitosan

To prepare BP-CS, water-soluble chitosan (CS, degree of deacetylation 95%, high molecular weight, Xi’an Lyphar Biotech) was dissolved in deionized (DI) water to create a solution with a concentration of 2% (w/v). The solution was stirred for 24 h at room temperature until fully dissolved, yielding a clear solution with no visible particulates. The pH of the solution ranged from 6.4 to 6.7.

The custom peptide WKTKTVPY (Biomatik) was dissolved in DI water at 2 mg/mL following the manufacturer’s recommendations. To activate peptide carboxyl groups, 1-ethyl-3-(3-dimethylaminopropyl) carbodiimide (EDC) and N-hydroxysuccinimide (NHS) were added to the peptide solution at a molar ratio of 3:1 relative to peptide carboxyl groups. This mixture was mixed for about 1 minute for activation and immediately added to the chitosan solution. The reaction proceeded for 24 hours under gentle stirring. For every 1 g of chitosan, 1 mg of peptide was used. Based on the feed ratio and assuming a typical EDC/NHS coupling efficiency (∼40%), the nominal grafting density is ∼0.39 μmol peptide g^−1^ chitosan. Following the reaction, the mixture was dialyzed against DI water using dialysis membrane with a molecular weight cutoff of 3500 Da for 24 hours, with three water changes to remove unreacted reagents and by-products. The dialyzed solution was lyophilized to obtain BP-CS as a dry powder.

### Fluorescent labelling of CS and BP-CS

Fluorescently labeled CS and BP-CS were prepared using fluorescein isothiocyanate (FITC). Chitosan (CS or BP-CS; 0.4 g) was dissolved in 40 mL of 80 mM acetic acid. FITC was dissolved in anhydrous methanol at 1 mg/mL and protected from light. A 1 M NaOH solution was prepared in DI water. For labeling, 20 mL of chitosan solution was mixed with an equal volume of anhydrous methanol and stirred for 3 h. The mixture was degassed under vacuum for 15 min, covered with aluminum foil, and FITC solution was added dropwise under continuous stirring. After reacting for 1 h at room temperature, 5 mL of 1 M NaOH was added to precipitate the chitosan. The precipitate was centrifuged at 4400 rpm for 5 min, washed with 30 mL DI water, and centrifuged again. The fluorescent chitosan (FITC-CS or FITC-BP-CS) was resuspended in DI water and dialyzed against DI water for 3 days, with water changes twice daily. The dialyzed solution was lyophilized.

### Hydrogel synthesis

An acrylamide precursor solution was prepared for both single-network PAAm gels and double-network Alg-PAAm tough gels. For PAAm gels, acrylamide was dissolved in water at 12% (w/v). For Alg– PAAm gels, acrylamide (12% w/v) and alginate (2% w/v) were dissolved in water. Precursor solutions were stirred until fully dissolved (30 min for PAAm; 48 h for Alg–PAAm). Prior to polymerization, solutions were degassed for 5 min. Ammonium persulfate (APS) was prepared at 66 mg/mL in water. N,N’-methylenebisacrylamide (MBAA) was prepared as a 2% (w/v) solution in water. Calcium sulfate slurry was prepared at 0.75 M and stirred continuously for 30 min before use.

For PAAm gels, precursor solution was mixed with 64.8 µL MBAA solution and 32 µL TEMED, then syringe-mixed with 452 µL APS solution and 250 µL water. The mixture was immediately injected into a closed glass mold and polymerized overnight. The Alg-PAAm hydrogel (pH = 7) was prepared following a previously reported protoco**l** ^**35**^. The precursor solution was mixed with 72 µL MBAA solution, 32 µL TEMED, and then syringe-mixed with 452 µL APS solution, and 250 µL calcium sulfate slurry before molding and overnight polymerization. Hydrogels were stored in humid environment at 4 °C before use.

### FTIR and NMR characterization

CS and BP-CS were characterized using Fourier-transform infrared spectroscopy (FTIR) and nuclear magnetic resonance spectroscopy (NMR) to confirm peptide conjugation and analyze structural modifications. FTIR analysis was conducted using a Perkin Elmer Spectrum Two FT-IR Spectrometer over 4000–500 cm^−1^ with a resolution of 1 cm^−1^. Lyophilized samples were ground into powders prior to analysis. Characteristic peaks, including those corresponding to amide bond formation, such as C=O stretching (amide I band) and N-H bending (amide II band), were examined. ^1^H NMR spectra were collected using a Bruker AVIIIHD 500 MHz spectrometer. Samples were dissolved in deuterium oxide (D2O) at a concentration of 50mg/mL. Signals corresponding to the chitosan backbone were identified, while additional aromatic and aliphatic peaks associated with the peptide sequence were examined in BP-CS.

### Peeling tests

Porcine large intestinal tissue was gently rinsed with PBS. At room temperature, tissue was cut into approximately 1.5cm wide segments, unrolled, and trimmed to 1.5 cm by 6 cm specimens. Samples were kept humid prior to use.

For adhesion energy measurements, 0.5 mL of CS or BP-CS solution was evenly applied to tissue surfaces, followed by placement of PAAm or Alg–PAAm hydrogels. T-peel tests were conducted using an Instron machine (Model 5965) at a peeling rate of 60 mm/min. Adhesion energy was calculated using the formula 2*F*_avg_/W, where *F*_*avg*_ is the steady-state peeling force and W is sample width.

### Mucin binding assay

The mucin binding assay was adapted from published protocols ^31^. ELISA 96-well plates were coated with porcine gastric mucin type III (10 µg/mL; Sigma-Aldrich) and incubated overnight at 4°C in 50μL carbonate-bicarbonate buffer (pH 9.6). Plates were blocked with 1% bovine serum albumin (BSA) in PBS for 1 hour, then incubated with 0.2% chitosan solution for 1 hour at room temperature. Bound chitosan in each well was visualized using confocal microscope (Zeiss, LSM800), and fluorescence intensity was quantified using ImageJ.

### In vitro cytocompatibility assessment

Human vocal fold fibroblast (HVFF) cells were cultured in DMEM supplemented with 10% fetal bovine serum and 1% penicillin-streptomycin. Cells were maintained at 37°C in a humidified atmosphere containing 5% CO2. Prior to cytotoxicity tests, PAAm and Alg-PAAm hydrogels underwent solvent exchange in DI water (three 12-h exchange) to remove unreactive precursors and sterilized with UV light exposure for at least 1 hour. Extracts were prepared by incubating 200 mg of hydrogel in 1 mL DMEM for 24 h. Cells were seeded into 96-well plates at 50,000 cells per well. After 24 hours, culture medium was replaced with extract media supplemented with 1% penicillin-streptomycin and 10% fetal bovine serum. Pristine cell culture medium was used as control. Live/Dead staining (Invitrogen) was performed after 24 h according to the manufacturer’s protocol, and samples were imaged using confocal microscopy.

### Intestinal drug diffusion study

To prepared BSA-loaded hydrogels, FITC–BSA (0.5% w/v) was incorporated into precursor solutions prior to gelation. Gels were adhered to porcine intestinal tissue using BP-CS or CS with EDC/NHS and incubated for 6 h at room temperature. After carefully removing the gels, tissue samples were fixed in 4% paraformaldehyde in PBS for 24 hours. After fixing, samples were embedded in OCT, flash frozen, and then cryosectioned for histological analysis. Tissue sections were stained with Hoechst 33342 (H3570, Invitrogen) dye for nuclei visualization and wheat germ agglutinin conjugate (WGA) with Alexa Fluor 647 (W32466, ThermoFisher Scientific) for mucin staining. Slides were stained with Hoescht 33342 for 30min and washed thrice in PBS for 10 min. Mucin was stained with WGA according to supplier’s recommendation, then washed for 10 min. Confocal microscopy (Zeiss, LSM800) was used to quantify diffusion depth based on FITC fluorescence intensity.

### In vitro drug release characterization

FITC–BSA-loaded PAAm gels were placed in Transwell inserts (9310412, cellQART®) with CS, BP-CS, or CS + EDC/NHS interfaces and incubated at 37 °C, allowing diffusion of BSA-FITC into 200 µL PBS in 12 well plates. Release medium was collected at specified time points (0.5, 1, 2, 4, 12, 24 hours) and analyzed using a fluorescence plate reader (Tecan Infinite® 200 Pro). Drug concentration was calculated using a standard calibration curve.

### Statistical analysis

All experiments were performed with a minimum sample size of n ≥3 Data are reported as mean ± standard deviation (SD). Statistical comparisons between two groups were conducted using independent samples t-tests. P-values < 0.05 were considered statistically significant, while **\*\*\* corresponds with P-values < 0.001, ** with P-values < 0.01, and * with P-values < 0.05**. Statistical analyses were performed using OriginLab and GraphPad Prism software.

## Author contributions

Evan Johnston contributed to conceptualization, methodology, formal analysis, investigation, writing – original draft, writing – review and editing, visualization. Megan Farrow contributed to data curation, formal analysis, investigation, methodology, validation, writing – original draft, review, editing, visualization. Zhen yang contributed to formal analysis, methodology, writing – review and editing. Aram Bahmani contributed to methodology, writing – review and editing. Yin Liu contributed to methodology, writing – review and editing. Xin Huang contributed to conceptualization, writing – review and editing. Jing Yan contributed to conceptualization, writing – original draft, review, editing. Jianyu Li contributed to conceptualization, writing – original draft, review, editing, visualization, supervision, project administration, funding acquisition.

## Conflicts of interest

There are no conflicts to declare.

## Data availability

The data supporting this article, including NMR spectra and frequency sweep rheology have been included as part of the supplementary information (SI). Supplementary information is available. See DOI: []

## Acknowledgements

This work is supported by New Frontiers in Research Fund (NFRFE-2022-00348), Natural Sciences and Engineering Research Council of Canada (RGPIN-2024-04925), National Institutes of Health of the United States (R01DC021461). J.L. acknowledge financial support from the Canada Research Chairs Program. J.Y. acknowledges financial support from the Burroughs Wellcome Fund (#1022835).

## Notes

### Competing Interest Statement

The authors have declared no competing interest.

### Summary of Updates

Only last name of fist author has been changed from Hohnston to Johnston.

