## Supplemental for "Bacteria-mimetic bioadhesives with multivalent mucoadhesion and drug-compatible delivery"

Electronic Supplemental Information (ESI) for:

**Figures**

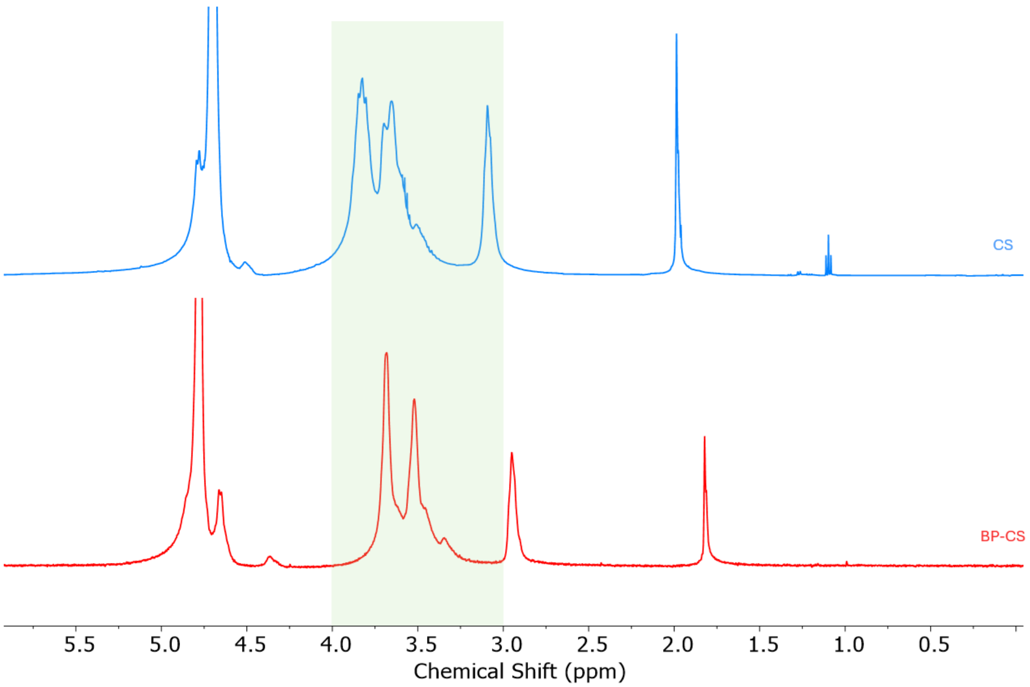


**Fig. S1 NMR valuation and confirmation of chitosan modification with bacterial peptide.** NMR Spectra for pristine chitosan (CS) and bacterial peptide modified chitosan (BP-CS). Signal shifts around 3 – 4 ppm suggest aliphatic side chains or modifications to the CS backbone caused by peptide attachment, validating the modification.

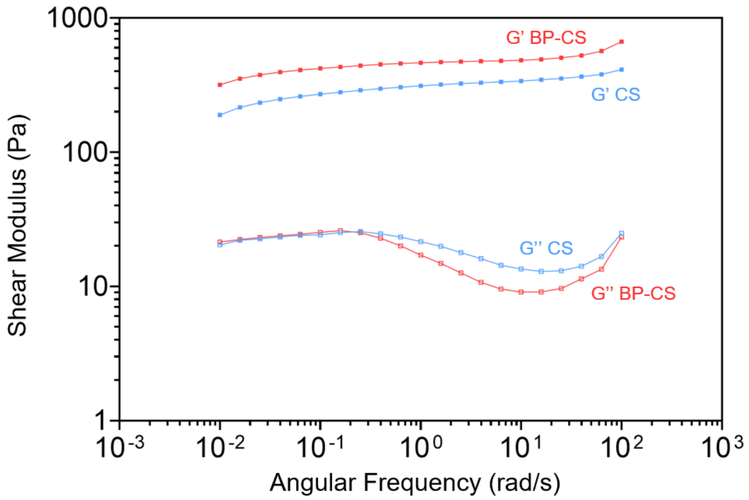


**Fig. S2 Frequency sweep mechanical response.** Frequency sweep mechanical response of pristine chitosan (CS) and bacteria-peptide modified chitosan (BP-CS), showing comparable storage modulus (G') and loss modulus (G'') over a wide range of angular frequencies. This result further suggests that the grafting of the short Bap1-derived peptide motif on chitosan does not result in alterations to its inherent mechanical properties
